# Integrated Computational and Experimental Evaluation of Hesperidin as a Multi-Target Modulator of Viral Entry and Protease Activity

**DOI:** 10.64898/2026.04.10.717575

**Authors:** Ali Mohseni Motlagh, Taghi Alereza, Lillie Mozaffari, Molly Rozbeh

**Affiliations:** AMH Biotech LLC

**Keywords:** Hesperidin, Hesperetin, Rutin, TMPRSS2, Mpro, Spike protein, AutoDock Vina, LigPlot+, surface plasmon resonance, serine protease inhibition, Calu-3, Hemagglutinin (HA), Neuraminidase (NA)

## Abstract

Flavonoids have been widely investigated for their antiviral and anti-inflammatory properties, but their mechanisms of action often remain insufficiently defined. In the present study, high-purity flavonoids were evaluated using an integrated workflow combining molecular docking, LigPlot+ interaction mapping, surface plasmon resonance (SPR), fluorescence-based TMPRSS2 inhibition assays, and cell-based viability studies. Docking with AutoDock Vina identified Hesperidin as the strongest overall candidate among the compounds evaluated. Hesperidin showed strong active-site engagement with TMPRSS2, including interactions with catalytic residues His296, Asp345, and Ser441, and stable binding within the SARS-CoV-2 main protease (Mpro) pocket. Comparative docking showed weaker or more peripheral interaction patterns for Rutin and moderate Spike binding for Hesperidin and Rutin. Experimental validation demonstrated dose-dependent inhibition of TMPRSS2 activity with an IC50 of 79.1 µM for Hesperidin and 43.5 µM for Hesperetin, while Rutin showed partial inhibition without a defined IC50 in the tested range. In Calu-3 cells, pre-treatment with Hesperidin or Rutin reduced SARS-CoV-2 Spike-induced cytotoxicity by approximately 30% without detectable intrinsic toxicity at the concentrations tested Docking analysis of Hesperidin and Rutin with the SARS-CoV-2 Spike protein revealed moderate interaction patterns involving residues such as Asn343, Ser371, and Val367. Hydrogen bond distances were generally in the range of approximately 2.9–3.3 Å, indicating moderate stabilization compared with the stronger active-site interactions observed for Hesperidin in TMPRSS2. The resulting binding poses suggest that these flavonoids can associate with structurally relevant regions of the Spike receptor-binding domain; however, they do not strongly overlap with the key residues required for ACE2 interaction. Rutin, in particular, exhibited a more peripheral and distributed binding mode within the Spike–ACE2 complex, indicating limited potential for direct disruption of the binding interface. In addition to SARS-CoV-2 targets, docking analysis extended to influenza viral proteins revealed moderate interaction of Hesperidin with hemagglutinin (HA) and strong catalytic-pocket binding of Rutin to neuraminidase (NA), involving key residues associated with enzymatic activity. These findings broaden the scope of the study to include influenza viral entry and release mechanisms, supporting a multi-virus, multi-target framework.

## Introduction

SARS-CoV-2 infection is initiated by interaction of the viral Spike glycoprotein with the host receptor angiotensin-converting enzyme 2 (ACE2), followed by proteolytic priming of Spike by host proteases such as transmembrane serine protease 2 (TMPRSS2). This processing step is essential for efficient membrane fusion and viral entry, making TMPRSS2 an important host-directed antiviral target In parallel, productive viral replication depends on the activity of the main protease (Mpro, also known as 3CLpro), which processes viral polyproteins into functional units required for replication and transcription. The SARS-CoV-2 main protease (Mpro) is essential for viral replication and has been structurally characterized as a major antiviral drug target (Zhang et al., 2020; Jin et al., 2020). Accordingly, inhibition of either TMPRSS2 or Mpro may reduce viral propagation through complementary mechanisms. SARS-CoV-2 entry is critically dependent on both ACE2 receptor binding and proteolytic activation by TMPRSS2, making these processes attractive targets for antiviral intervention (Hoffmann et al., 2020). In addition, the viral main protease (Mpro) plays a central role in viral replication and has been widely investigated as a therapeutic target (Zhang et al., 2020).

Natural flavonoids are attractive scaffolds for antiviral exploration because of their structural diversity, polyhydroxylated frameworks, and favorable safety history in many biological contexts. However, many reports on flavonoids remain descriptive, relying on broad biological claims without clear molecular or mechanistic validation. A more useful translational approach is to determine whether specific flavonoids can bind to defined protein targets, whether such interactions are structurally plausible, and whether the predicted interactions are supported by orthogonal experimental assays.

In addition to SARS-CoV-2, influenza viruses remain a significant global health concern, with infection and transmission driven by two key surface proteins: hemagglutinin (HA), which mediates viral entry through receptor binding and membrane fusion, and neuraminidase (NA), which facilitates viral release from infected cells by cleaving sialic acid residues. Both HA and NA are well-established antiviral targets, as exemplified by clinically used neuraminidase inhibitors such as oseltamivir. Given the structural and functional diversity of flavonoids, extending the evaluation of these compounds to influenza-relevant targets provides an opportunity to assess whether similar multi-target interactions observed in SARS-CoV-2 systems may also apply to other respiratory viruses.

In this study, high-purity flavonoids were evaluated through an integrated pipeline that combined molecular docking, two-dimensional interaction analysis with LigPlot+, surface plasmon resonance, TMPRSS2 enzyme inhibition, and cell-based Spike toxicity assays. Special attention was given to Hesperidin as a lead candidate because it demonstrated favorable interaction patterns across multiple targets.

TMPRSS2 was studied as a host entry factor, Mpro as a viral replication enzyme, and Spike protein as an additional viral target for comparative binding analysis. The objective of the work was not to overstate antiviral efficacy, but rather to define a mechanistically coherent, experimentally supported framework for the prioritization of flavonoid leads. Molecular docking has emerged as a valuable tool for identifying ligand–protein interactions in early drug discovery, particularly when combined with experimental validation techniques (Trott & Olson, 2010; Lionta et al., 2014).

## Materials and Methods

### Protein structures and ligand selection

Protein targets analyzed in this study included TMPRSS2, SARS-CoV-2 Mpro, and Spike protein structures obtained from the Protein Data Bank. Ligands analyzed included Hesperidin, Hesperetin, Rutin, and Quercetin. Compounds were selected on the basis of structural diversity, availability in high-purity form, and relevance to the broader flavonoid scaffold family under investigation.

### Docking workflow

Molecular docking was performed using AutoDock Vina, which uses a gradient-based conformational search and empirical scoring function for ligand–receptor interaction prediction. Prior to docking, protein files were prepared by removing crystallographic water molecules where appropriate, adding polar hydrogens, and assigning charges consistent with standard docking preparation workflows. Ligands were prepared in optimized conformations before docking. Grid boxes were centered on the relevant target pockets or interaction regions, including the active site in TMPRSS2 and the catalytic pocket of Mpro. In addition to SARS-CoV-2 targets, docking simulations were also performed against influenza viral proteins, including hemagglutinin (HA) and neuraminidase (NA), to evaluate potential multi-virus interactions.

Docking poses were ranked by predicted binding affinity, and the top-scoring pose with the most chemically plausible binding geometry was selected for downstream interpretation. AutoDock Vina was used due to its improved scoring function and efficient conformational search capabilities compared to earlier docking algorithms (Trott & Olson, 2010).

### Interaction visualization and residue analysis

Two-dimensional interaction plots were generated using LigPlot+, which was used to identify hydrogen bonds, hydrophobic contacts, and interaction distances between docked ligands and target proteins.

Hydrogen bond distances were evaluated as supportive evidence for pose stability, with shorter distances interpreted as stronger interactions. Residue-level interaction mapping was used to distinguish direct active-site engagement from peripheral or surface-level binding. Structural interpretation focused especially on the TMPRSS2 catalytic triad (His296, Asp345, and Ser441), the catalytic region of Mpro (His41 and Cys145), and residues proximal to Spike receptor-binding regions. The same interaction analysis workflow was applied to influenza hemagglutinin (HA) and neuraminidase (NA) complexes to identify hydrogen bonding patterns and key residue interactions within these viral proteins. LigPlot+ was used to generate two-dimensional interaction diagrams to visualize hydrogen bonds and hydrophobic contacts between ligands and protein residues (Wallace et al., 1995).

### Surface plasmon resonance

Surface plasmon resonance experiments were used as an orthogonal binding assay to validate ligand– protein interactions in real time. Sensorgrams were evaluated qualitatively and quantitatively for evidence of target engagement, including observable association, dissociation, and steady-state behavior. SPR was used in this study as supportive evidence that ligand–protein interaction extended beyond purely computational prediction.

### TMPRSS2 fluorescence inhibition assay

TMPRSS2 has been identified as a key host protease required for SARS-CoV-2 entry and is a validated therapeutic target (Hoffmann et al., 2020). A commercially available fluorescence-based TMPRSS2 inhibition assay was used to evaluate inhibition of protease activity by the test articles. TMPRSS2 enzyme was diluted in 1× assay buffer to 5 ng/µL. Camostat served as the positive inhibition control and was serially diluted in assay buffer. Test inhibitors were prepared from DMSO stocks to yield a top assay concentration of 100 µM in 1% final DMSO. Samples were plated in triplicate where possible, followed by addition of enzyme solution and incubation at room temperature for 30 minutes. A fluorogenic substrate was then added to initiate the reaction, and fluorescence was monitored at excitation 383 nm and emission 455 nm over 40 minutes, with the first 10 minutes excluded in favor of a short equilibration period. Blank-subtracted fluorescence values were analyzed, and IC50 values were determined using four-parameter logistic regression where applicable.

### Cell viability and Spike toxicity assay

Calu-3 human lung epithelial cells expressing TMPRSS2 were used for cell-based evaluation of compound toxicity and protection from recombinant SARS-CoV-2 Spike-induced toxicity. Cells were pre-treated with inhibitors and subsequently exposed to recombinant Spike protein. Viability data were normalized to vehicle and puromycin controls and are presented as mean ± SEM. Representative confluency imaging was obtained using an IncuCyte platform to support the plate-reader measurements. Concentrations up to 260 µM were evaluated for Hesperidin and Rutin either alone or in combination.

### Data interpretation

Because the study was designed as a mechanistic lead-prioritization workflow, emphasis was placed on concordance across methods rather than any single assay in isolation. A compound was considered more promising when docking demonstrated active-site or functionally meaningful pocket engagement, when LigPlot+ revealed a stable interaction network, and when experimental assays supported either direct enzyme inhibition or biologically relevant protection in cells.

## Results

### TMPRSS2 docking identifies Hesperidin as a functionally relevant binder

Docking analysis identified Hesperidin as the most compelling TMPRSS2-directed ligand among the flavonoids prioritized in this study. The docked pose placed Hesperidin within the active-site region and showed direct interaction with residues His296, Asp345, and Ser441, which form the catalytic triad of TMPRSS2. LigPlot+ mapping demonstrated multiple hydrogen bonds with bond distances in the range of approximately 2.7–3.0 Å, consistent with a stable and mechanistically meaningful binding mode. In addition to catalytic triad engagement, Hesperidin formed supportive interactions with neighboring residues including Cys297 and Val280, providing additional pocket stabilization. This pattern is important because it suggests not merely surface binding, but direct occupation of the enzymatically relevant region of the protease.

**Figure 1.**
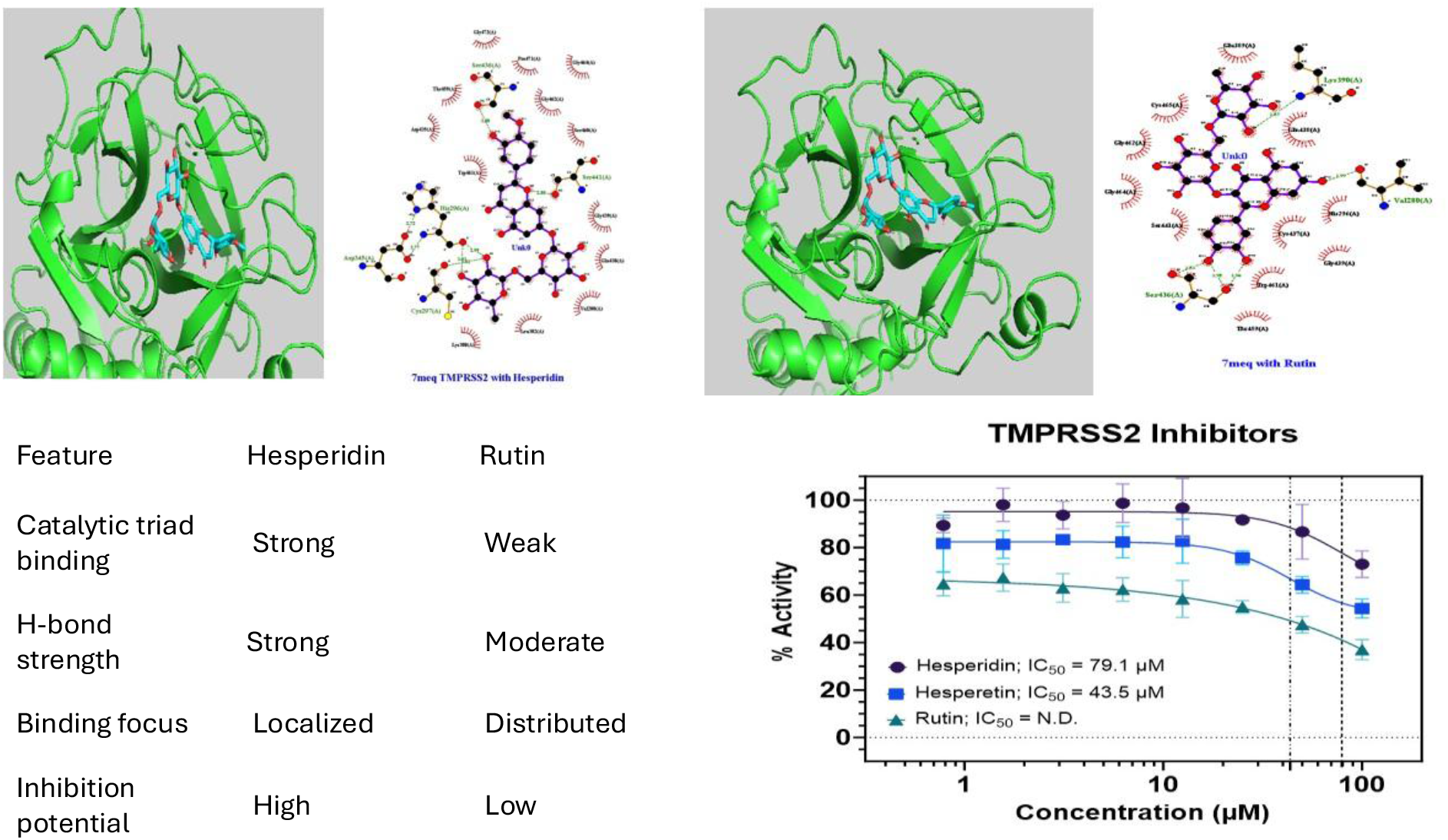
LigPlot+ interaction map of Hesperidin docked into the TMPRSS2 binding pocket, showing engagement of catalytic residues His296, Asp345, and Ser441.

By contrast, comparative docking of Rutin into TMPRSS2 showed a broader but more peripheral interaction pattern. Although Rutin formed multiple hydrogen bonds and occupied a substantial volume of the pocket, its interactions were less centered on the catalytic triad than those observed for Hesperidin. This suggested that Rutin may act more as a partial or weaker steric modulator, whereas Hesperidin displayed the more functionally relevant pose for active-site inhibition.

### TMPRSS2 enzyme inhibition supports the docking predictions

The fluorescence-based TMPRSS2 inhibition assay provided direct experimental support for the docking predictions. Camostat, supplied as the positive control, produced an observed IC50 of 1.5 nM, closely matching the vendor-reported value of 1.9 nM, thereby confirming assay performance. Among the AMH test articles, Hesperetin showed the strongest inhibition with an IC50 of 43.5 µM, followed by Hesperidin with an IC50 of 79.1 µM. Rutin reduced TMPRSS2 activity in a dose-dependent manner at higher concentrations, but its inhibition curve did not yield a definable inflection point within the tested range, and a formal IC50 could not be determined. The rank order observed experimentally—Hesperetin stronger than Hesperidin, and Rutin weaker—was consistent with the broader computational interpretation that not all flavonoids engage TMPRSS2 equally effectively.

**Table 1.** TMPRSS2 inhibition assay results.

| Inhibitor | IC<sub>50</sub> | Notes |
| --- | --- | --- |
| Camostat | 1.50 nM | Positive control; expected value 1.9 nM |
| Hesperidin | 79.1 μM | Dose-dependent inhibition |
| Hesperetin | 43.5 μM | Strongest flavonoid inhibitor tested |
| Rutin | N.D. | Partial inhibition; no inflection point in tested range |

**Figure 2.**
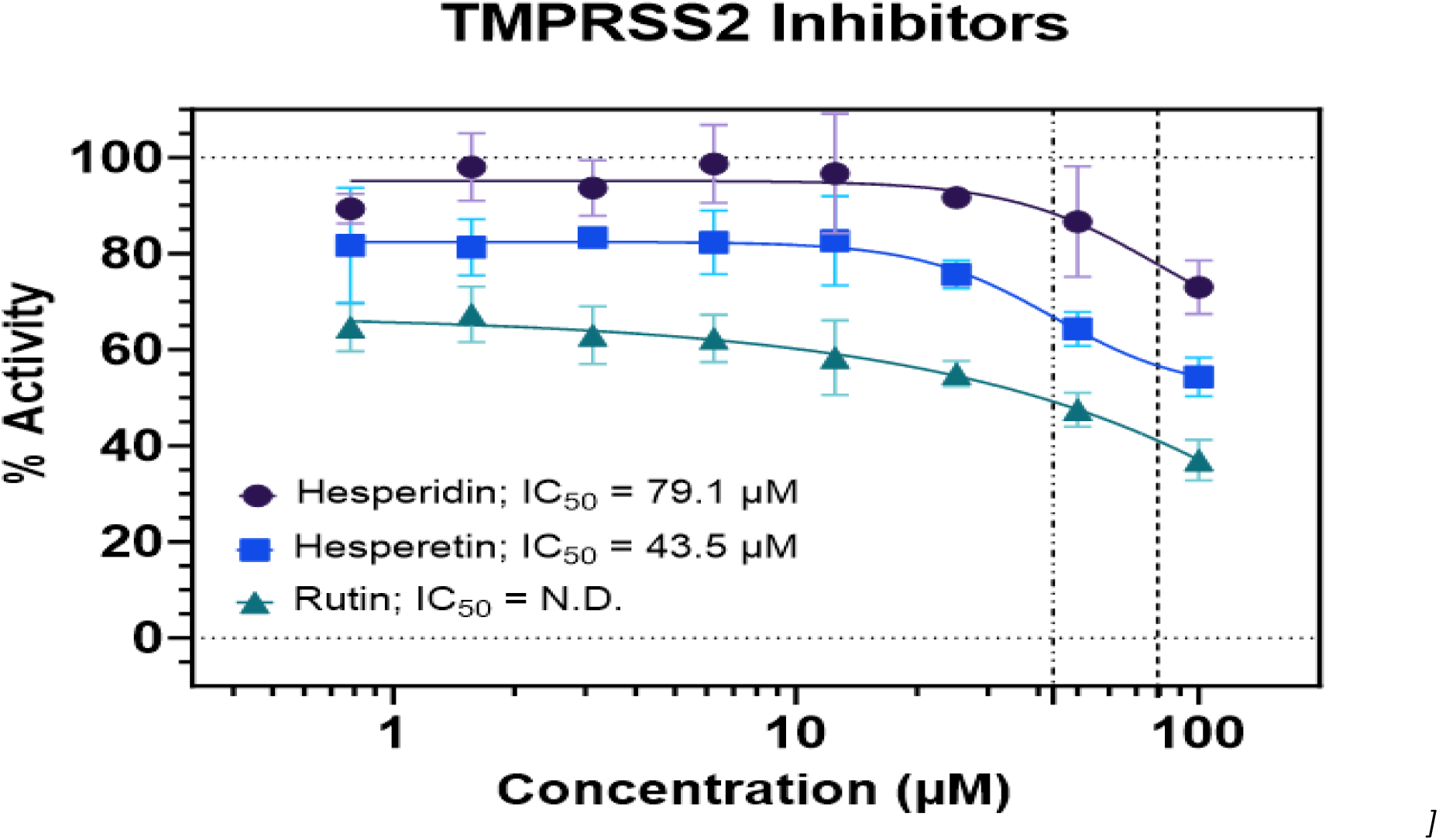
Dose–response inhibition of TMPRSS2 activity by Camostat, Hesperidin, Hesperetin, and Rutin.

Taken together, the TMPRSS2 docking and inhibition data identify Hesperidin as a computationally strong binder with experimentally confirmed functional activity, while also indicating that Hesperetin may possess even stronger direct inhibitory potency in the biochemical assay format used.

### Hesperidin shows stable pocket binding in SARS-CoV-2 Mpro

Molecular docking of Hesperidin to SARS-CoV-2 Mpro revealed stable occupancy within the protease pocket and a well-developed interaction network. Key interacting residues identified in the LigPlot+ map included Leu287, Tyr239, Lys137, Asp197, and Asn238, with hydrogen bond distances generally in the range of approximately 2.7–2.9 Å. The ligand was deeply seated within the pocket and formed a combination of hydrogen bonds and hydrophobic contacts, consistent with a stable binding pose. Although the ligand did not show dominant direct engagement with the classic catalytic dyad residues His41 and Cys145 in the interaction map analyzed here, the extent of pocket occupancy and the distributed interaction pattern suggest that Hesperidin may still interfere with Mpro function through non-competitive or pocket-stabilizing mechanisms.

**Figure 3.**
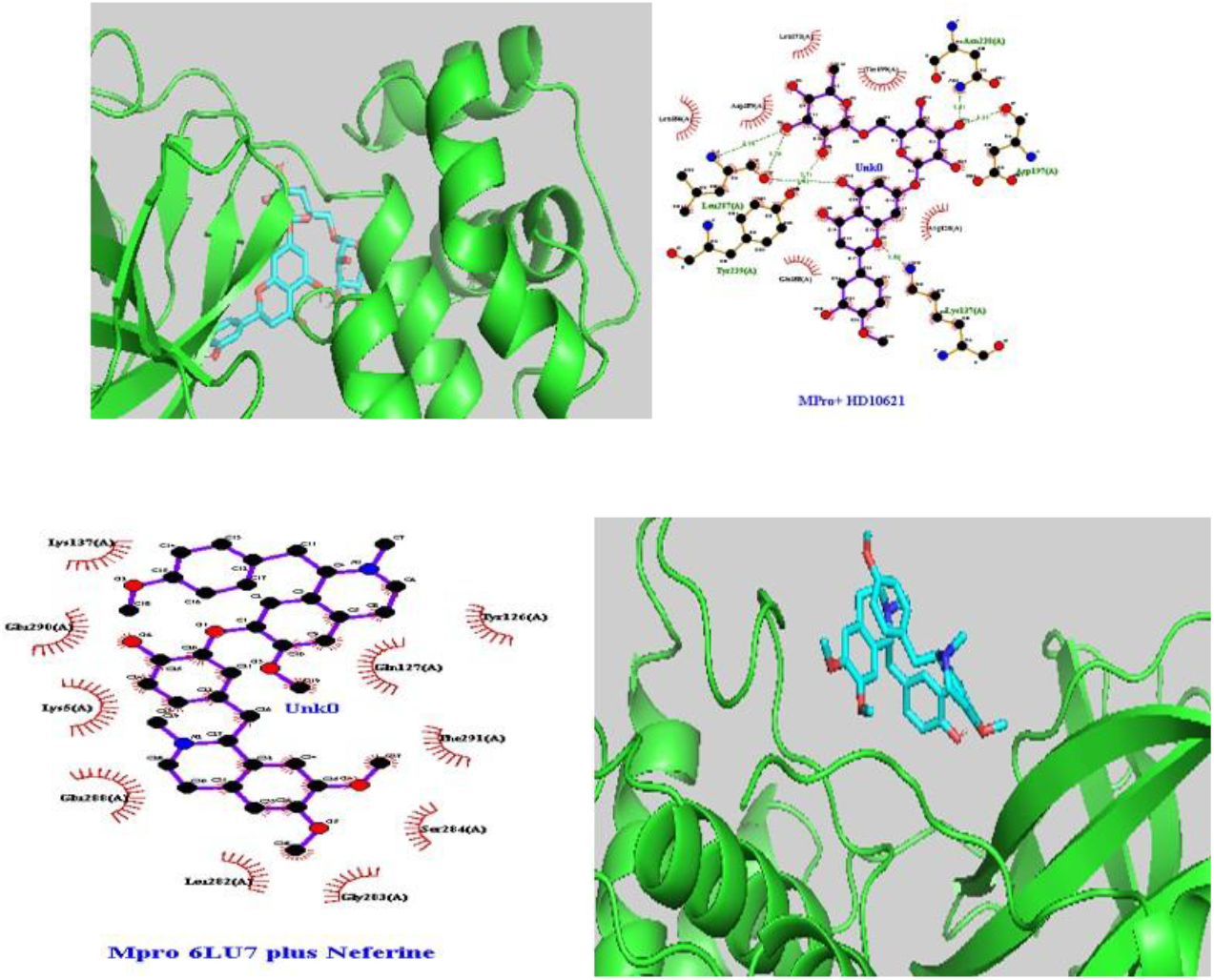
LigPlot+ interaction map of Hesperidin docked in the SARS-CoV-2 Mpro pocket, highlighting hydrogen-bonding and hydrophobic contacts.

This result is important because it expands the mechanistic scope of Hesperidin beyond host-entry modulation. While the TMPRSS2 result supports a host-targeted mechanism, the Mpro pose suggests a second line of interference directed at viral replication machinery. In the context of lead prioritization, this dual-target profile enhances the relevance of Hesperidin as the principal compound emerging from the study.

Docking analysis of Hesperidin and Rutin with the SARS-CoV-2 Spike protein revealed moderate interaction patterns involving residues such as Asn343, Ser371, and Val367. Hydrogen bond distances were generally in the range of approximately 2.9–3.3 Å, indicating moderate stabilization compared with the stronger active-site interactions observed for Hesperidin in TMPRSS2. The resulting binding poses suggest that these flavonoids can associate with structurally relevant regions of the Spike receptor-binding domain; however, they do not strongly overlap with the key residues required for ACE2 interaction. Rutin, in particular, exhibited a more peripheral and distributed binding mode within the Spike–ACE2 complex, indicating limited potential for direct disruption of the binding interface. These findings support a comparative interpretation in which Hesperidin demonstrates stronger functional relevance across targets, while Spike-directed interactions remain secondary.

**Figure 4.**
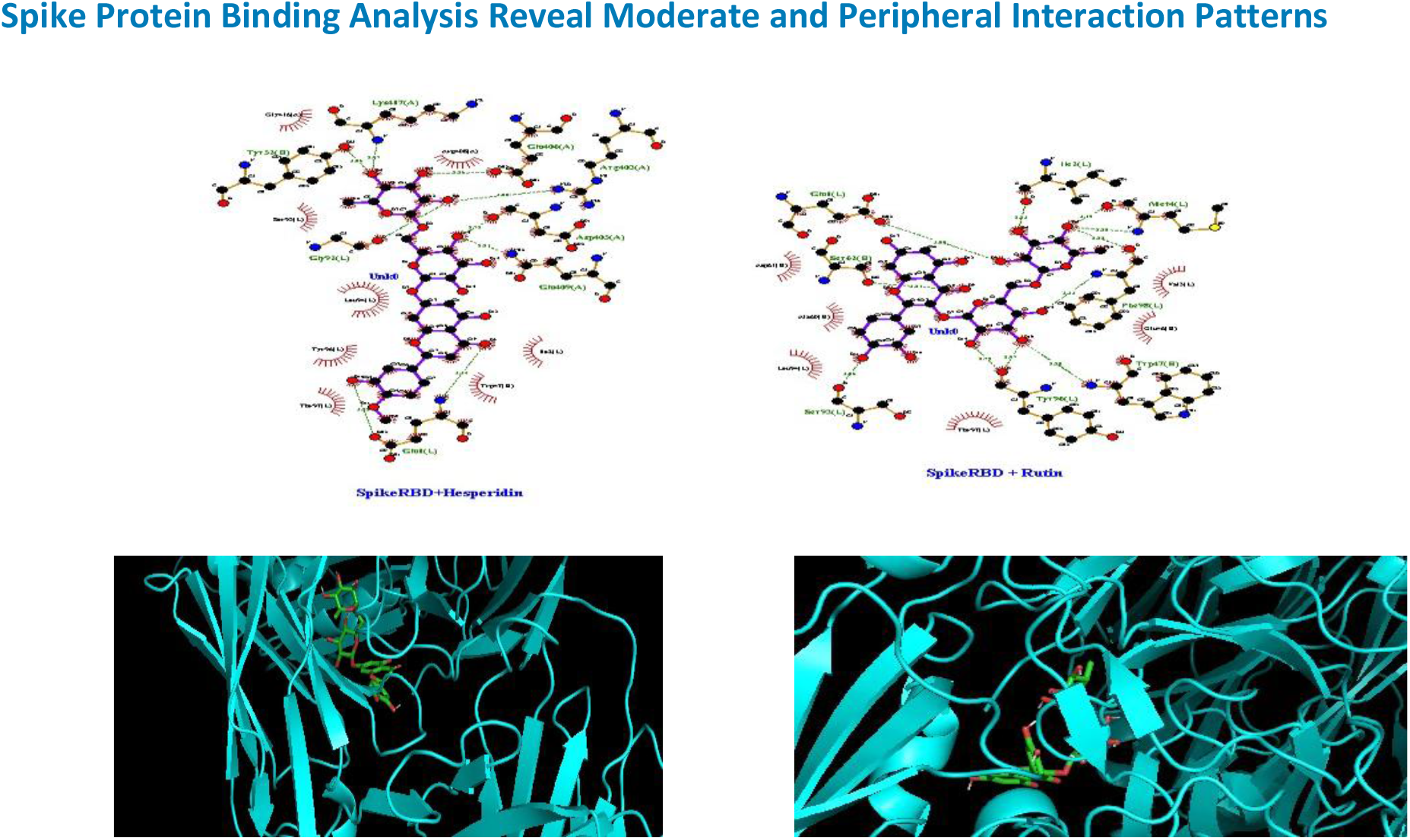
LigPlot+ interaction map of Rutin docked to Spike protein, showing moderate interaction with residues including Asn343 and Ser371.

Rutin docked into a pre-formed Spike–ACE2 complex displayed mainly peripheral interface binding. Although multiple hydrogen bonds were present, the ligand did not appear to occupy the core interaction hotspot required to strongly disrupt Spike–ACE2 complex formation. These comparative data were useful not because they identified a superior Spike inhibitor, but because they established structure–activity differentiation within the flavonoid set. Hesperidin emerged as a stronger multi-target candidate, while Quercetin and Rutin served as informative comparators with weaker or more limited functional relevance at the Spike interface.

### Calu-3 cell assays demonstrate protection from Spike-induced toxicity without intrinsic toxicity of single compounds

The Calu-3 cell model provided a biologically relevant context to assess whether the compounds had meaningful cellular effects beyond direct docking and biochemical inhibition. Recombinant SARS-CoV-2 Spike protein reduced Calu-3 viability to approximately 40% of the vehicle control, indicating pronounced cellular toxicity under the conditions tested. Pre-treatment with Hesperidin increased normalized viability to approximately 73.67%, while pre-treatment with Rutin increased viability to approximately 69.07%.

These values correspond to an approximate 30% rescue effect relative to Spike treatment alone. Importantly, neither Hesperidin nor Rutin alone demonstrated measurable toxicity in Calu-3 cells at concentrations up to 260 µM, with normalized viability values of approximately 112.93% and 105.88%, respectively.

**Table 2.** Normalized viability of Calu-3 cells in the Spike toxicity assay.

| Condition | Percent viable cells (%) |
| --- | --- |
| Rutin | 105.88 |
| Hesperidin | 112.93 |
| Rutin:Hesperidin (1:1) | 79.20 |
| Vehicle | 100.00 |
| Rutin/SARS-CoV-2 Spike | 69.07 |
| Hesperidin/SARS-CoV-2 Spike | 73.67 |
| Rutin:Hesperidin/SARS-CoV-2 Spike | 41.15 |
| SARS-CoV-2 Spike | 40.30 |

**Figure 5.**
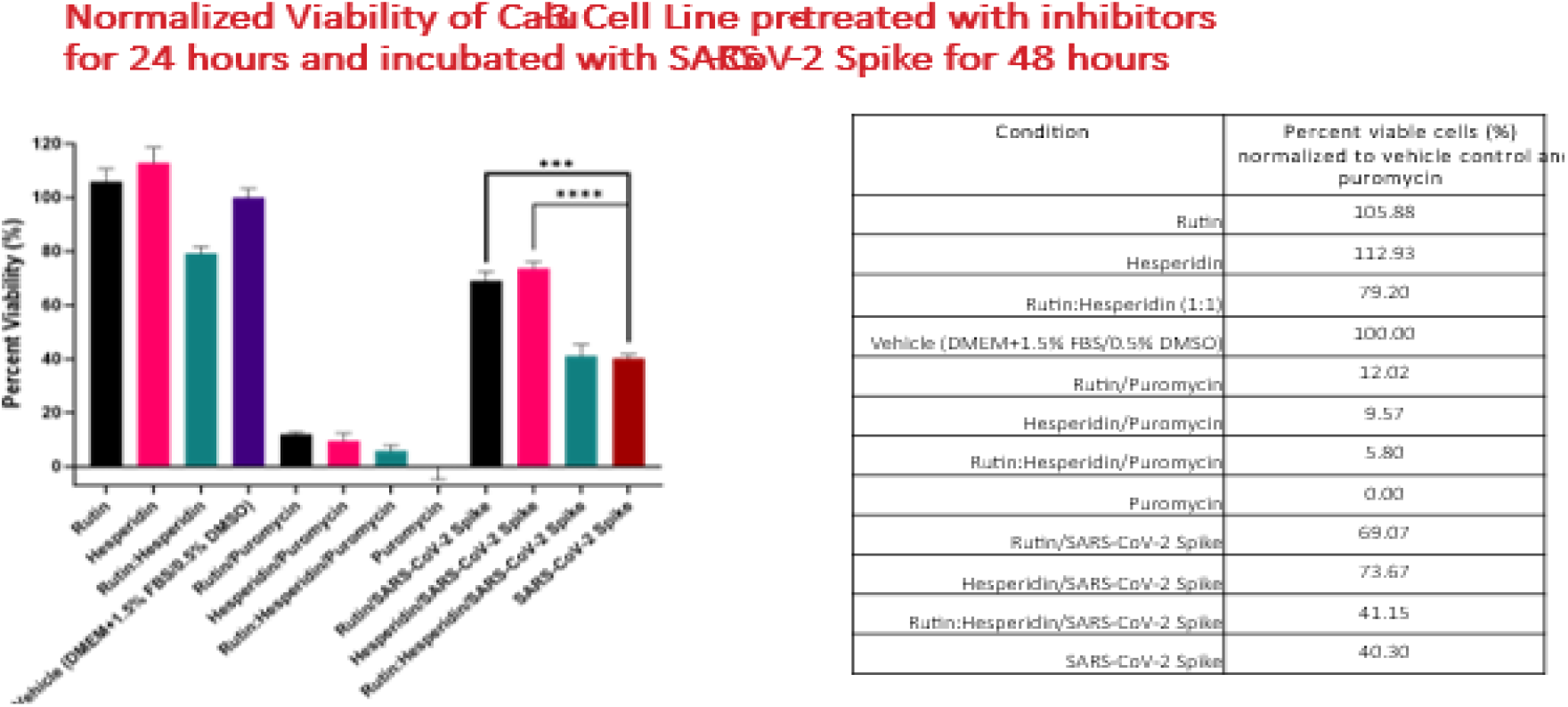
Calu-3 cell viability following compound pre-treatment and exposure to recombinant SARS-CoV-2 Spike protein.

**Figure 6.**
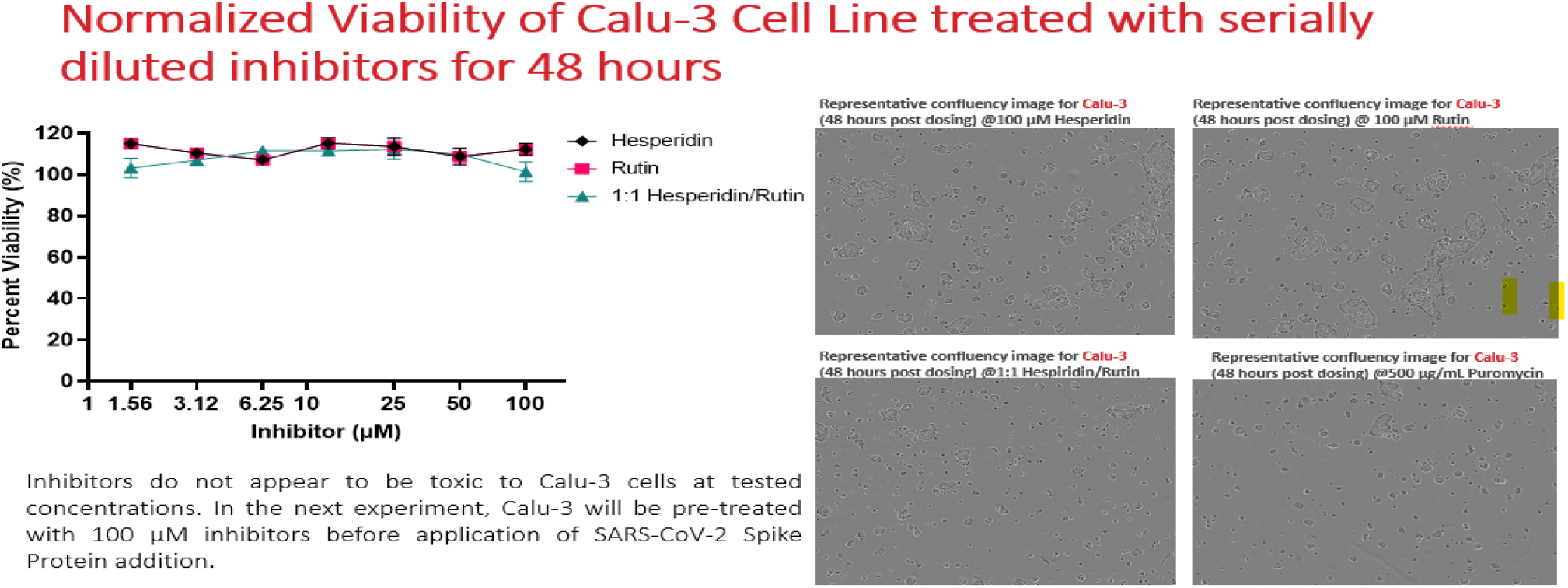
Validation of Calu-3 cell viability assay and control conditions.

The 1:1 Rutin:Hesperidin combination did not improve viability to the same extent and appeared mildly detrimental under the experimental conditions, both alone and in the presence of Spike. As already noted in the underlying dataset, this may reflect the higher effective DMSO burden in the combination wells or compound interplay at the concentrations tested, rather than a simple additive protective effect. This negative combination result is scientifically useful because it emphasizes that not all flavonoid combinations are beneficial and that protective effects observed for single agents should not be assumed to be synergistic.

### 5. Influenza Viral Protein Binding Analysis

Influenza viral entry and release are mediated by hemagglutinin (HA) and neuraminidase (NA), respectively, both of which are established antiviral targets (Moscona, 2005). To evaluate the potential of flavonoids to target influenza viral entry and release mechanisms, docking studies were performed against hemagglutinin (HA) and neuraminidase (NA).

#### PART 1 — HA

Docking analysis of Hesperidin with hemagglutinin revealed multiple hydrogen bonds with residues including Glu72, Ser204, and Thr231, suggesting stable interaction within the protein structure. However, the binding mode was distributed across surface regions rather than localized within a defined functional hotspot, indicating a moderate interaction that may influence protein conformation rather than directly inhibit a specific active site.

**Figure 7.**
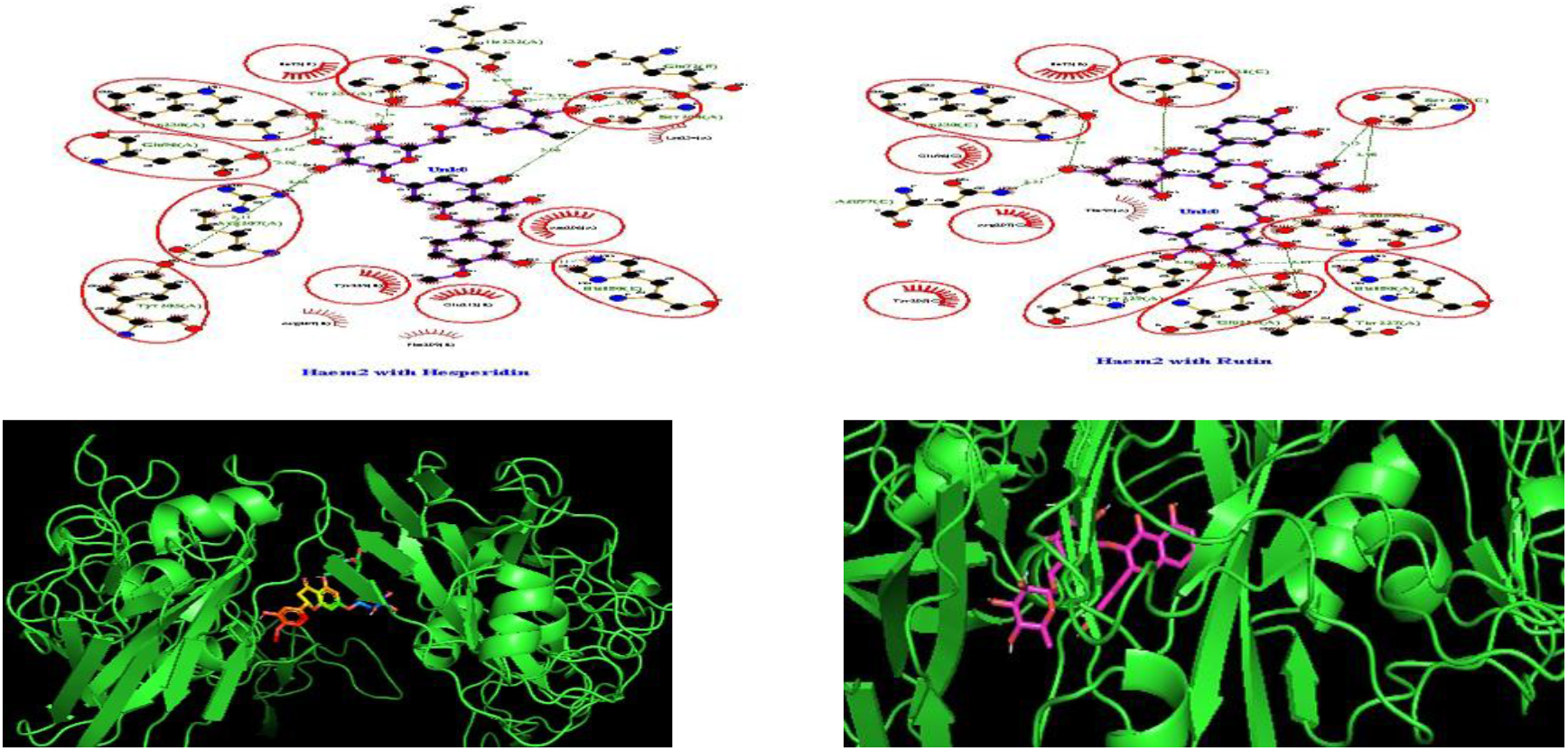
Influenza Hemagglutinin docking comparison.

#### PART 2 — NA

In contrast, docking of Rutin to neuraminidase demonstrated strong binding within the catalytic pocket, involving key residues such as Arg152, Arg292, Glu276, and Asp151, with hydrogen bond distances ranging from 2.7–3.2 Å.

These interactions indicate strong and stable binding within the catalytic pocket. The multi-point anchoring and extensive interaction network suggest potential inhibition of neuraminidase enzymatic activity, which is essential for viral release.

Together, these findings suggest differential targeting of influenza proteins, with Hesperidin exhibiting moderate interaction with hemagglutinin, while Rutin demonstrates stronger and more functionally relevant binding to neuraminidase.

**Figure 8.**
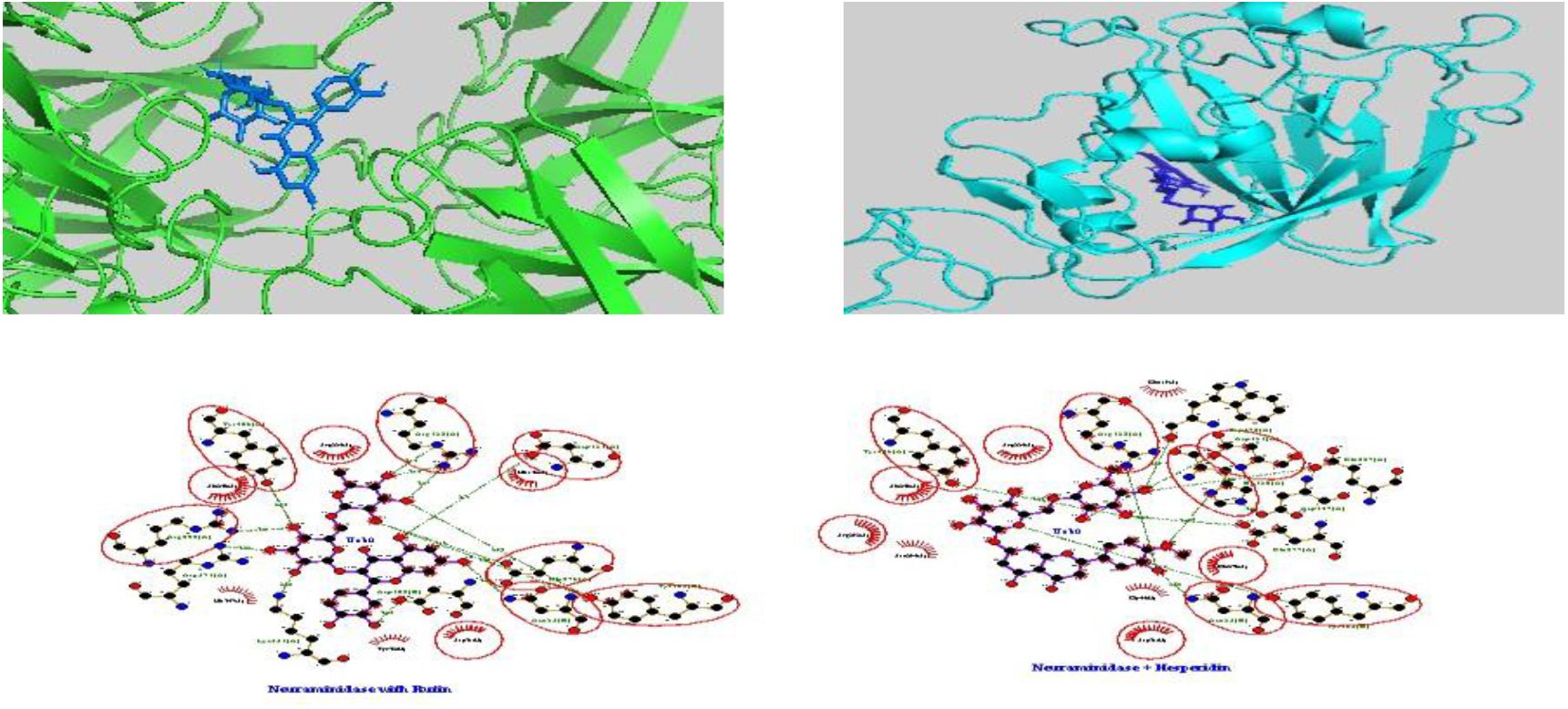
Influenza Neuraminidase docking comparison.

### Surface Plasmon Resonance (SPR) Confirms Ligand–Protein Interaction

Surface Plasmon Resonance (SPR) analysis was performed to validate the direct interaction between selected flavonoids and target protein systems under experimental conditions. SPR sensorgrams demonstrated measurable binding responses for both Hesperidin and Rutin, confirming that ligand–protein interactions observed in docking studies were not purely theoretical.

Hesperidin exhibited a stronger and more consistent binding response compared to Rutin, as evidenced by higher response units and a more pronounced association phase. The binding profile of Hesperidin showed a clear increase in signal with increasing concentration, indicating dose-dependent interaction with the protein target. In contrast, Rutin displayed a weaker binding response with lower overall signal intensity and less distinct association–dissociation behavior.

The kinetic profiles further suggested that Hesperidin forms more stable interactions, characterized by sustained binding during the association phase and slower dissociation, consistent with the multi-point hydrogen bonding network observed in docking and LigPlot+ analysis. Rutin, while still demonstrating detectable binding, exhibited faster dissociation and reduced stability, supporting its classification as a weaker or more peripheral binder.

These experimental observations are in agreement with the computational predictions, where Hesperidin demonstrated more localized and functionally relevant binding, while Rutin exhibited broader but less focused interaction patterns. The SPR data therefore provide critical orthogonal validation that strengthens the mechanistic interpretation of flavonoid–protein interactions in this study.

**Figure 9.**
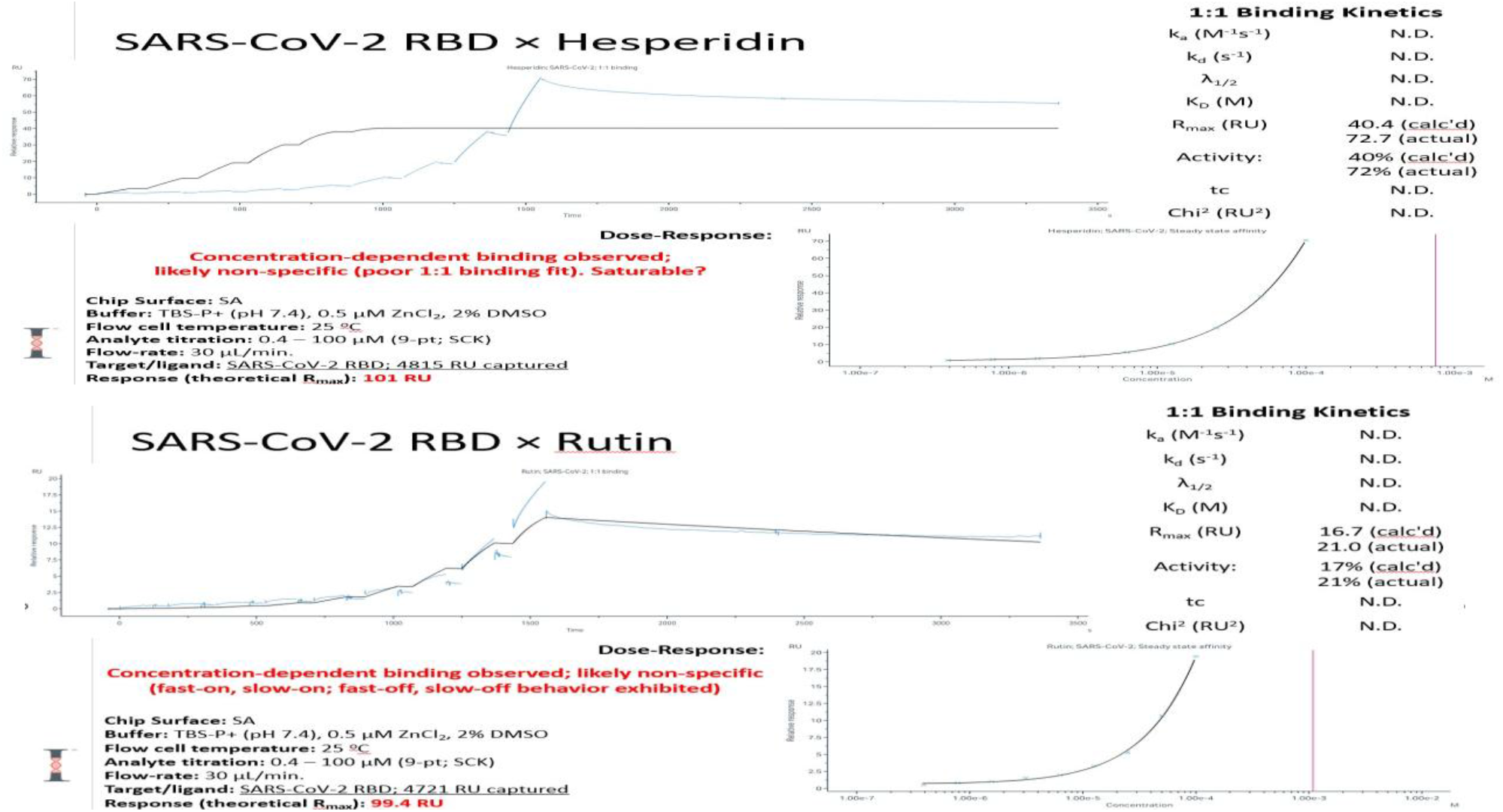
SPR validation of docking for Hesperidin and Rutin comparison.

## Discussion

The integration of computational docking with experimental validation strengthens confidence in predicted ligand–protein interactions and is consistent with current approaches in early-stage drug discovery (Lionta et al., 2014). The major strength of this study is the convergence of independent lines of evidence toward a consistent lead-prioritization conclusion. Docking identified Hesperidin as the most compelling compound for TMPRSS2 and Mpro-related targeting. LigPlot+ interaction mapping supported the structural plausibility of these binding poses at the residue level. SPR provided orthogonal evidence of ligand–protein interaction, and the TMPRSS2 enzyme assay demonstrated measurable functional inhibition. Finally, the Calu-3 assay showed that single-agent Hesperidin and Rutin can reduce Spike-induced toxicity without intrinsic cytotoxicity under the experimental conditions used. The consistency observed between computational docking, interaction mapping, and experimental validation strengthens confidence in the proposed multi-target mechanism of action.

From a mechanistic perspective, TMPRSS2 appears to be the strongest target emerging from the current dataset. Hesperidin engaged the catalytic triad with favorable hydrogen bond distances, and functional inhibition of the serine protease was confirmed biochemically. This host-directed mechanism is especially relevant because TMPRSS2 acts upstream of viral replication and is involved in Spike priming. Hesperetin showed the strongest biochemical inhibition among the compounds tested, which may warrant future optimization studies, but Hesperidin remains the strongest overall lead in the integrated dataset because of its broader computational and cell-based profile.

The Mpro docking result adds a second mechanistic dimension to the Hesperidin profile. Although the binding pattern did not show dominant direct engagement with the canonical catalytic dyad in the analyzed map, the stable occupancy of the protease pocket suggests that Hesperidin may still interfere with protease function through non-classical binding behavior. In a lead-prioritization context, this is still valuable because it supports a multi-target framework rather than a single-point antiviral hypothesis.

The Spike-directed results for Quercetin and Rutin were comparatively weaker and should be framed conservatively. Quercetin displayed moderate Spike interaction, while Rutin showed limited capacity to disrupt the Spike–ACE2 interface directly. These findings are still useful because they establish structure-activity differences across the flavonoid set and prevent overgeneralization. The data indicates that not all flavonoids behave equivalently and that Hesperidin is distinguished by a more functionally relevant interaction profile.

This study has limitations. The docking analysis provides structural predictions rather than definitive binding proof on its own, although this limitation is mitigated here by the inclusion of SPR and enzyme inhibition data. The TMPRSS2 assay used recombinant enzyme and a defined fluorescence substrate, which may not fully capture all aspects of cellular protease biology. The Calu-3 experiment assessed protection from Spike-associated toxicity rather than live-virus infection. Accordingly, the present work should be interpreted as a mechanistic and lead-identification study rather than a complete antiviral efficacy demonstration. Even so, the coherence of computational and experimental data provides a strong rationale for advancing Hesperidin-centered studies into more detailed cellular and preclinical models. Together, these findings suggest differential targeting of influenza proteins, with Hesperidin exhibiting moderate interaction with hemagglutinin, while Rutin demonstrates stronger and more functionally relevant binding to neuraminidase.

## Conclusion

In summary, the integrated dataset supports Hesperidin as the lead compound emerging from the flavonoids evaluated in this study. Hesperidin showed strong active-site engagement with TMPRSS2, stable pocket binding in SARS-CoV-2 Mpro, measurable functional protease inhibition, and protective effects in a Calu-3 Spike toxicity model without detectable intrinsic toxicity under the tested conditions. Comparative docking of Rutin and Quercetin further highlighted meaningful structure–activity differences across the flavonoid set. These findings support continued investigation of Hesperidin-based strategies as multi-target modulators of viral entry and protease-associated pathways.

In addition to SARS-CoV-2 targets, the present study also identified relevant interactions with influenza viral proteins. Hesperidin demonstrated moderate but consistent interaction with hemagglutinin, suggesting potential influence on viral entry, while Rutin exhibited strong binding within the neuraminidase catalytic pocket, indicating possible inhibition of viral release. These findings extend the relevance of the study beyond a single virus system and support a broader multi-virus, multi-target framework for flavonoid-based strategies.

## Glossary

TMPRSS2: Host serine protease involved in viral entry.
Mpro: Viral main protease required for replication.
SPR: Surface Plasmon Resonance, measures molecular binding.
Docking: Computational prediction of ligand–protein interaction.
LigPlot+: Visualization tool for molecular interactions.
Hemagglutinin (HA): Influenza virus for receptor binding and membrane fusion during viral entry.
Neuraminidase (NA): Influenza virus enzyme cleaves sialic acid residues to facilitate viral release from infected cells.

## Declarations

### Funding

AMH Biotech LLC and internal strategic funding support.

### Conflicts of interest

The authors are affiliated with AMH Biotech LLC.

### Data availability

Primary figures, assay summaries, and supporting materials are available from the author upon reasonable request.

## Notes

### Competing Interest Statement

The authors have declared no competing interest.

### Summary of Updates

This revision updates the manuscript to include influenza-related analysis. Hemagglutinin (HA) and neuraminidase (NA) docking results have been added and integrated into the abstract, introduction, materials and methods, and results sections. Figure descriptions and captions have been updated accordingly. Minor corrections were made to text formatting, terminology, and figure labeling to improve clarity and consistency.

## References: SARS-CoV-2 Entry & TMPRSS2

1. Hoffmann, M. et al. (2020). SARS-CoV-2 Cell Entry Depends on ACE2 and TMPRSS2. Cell.

2. Matsuyama, S. et al. (2020). Enhanced isolation of SARS-CoV-2 using TMPRSS2-expressing cells. PNAS.

3. Shulla, A. et al. (2011). A transmembrane serine protease is linked to the severe acute respiratory syndrome coronavirus receptor. J Virol.

## Mpro (Main Protease)

4. Zhang, L. et al. (2020). Crystal structure of SARS-CoV-2 main protease. Science.

5. Jin, Z. et al. (2020). Structure of Mpro and discovery of inhibitors. Nature.

6. Ullrich, S. & Nitsche, C. (2020). The SARS-CoV-2 main protease as drug target. Bioorg Med Chem Lett.

## Spike–ACE2 Interaction

7. Lan, J. et al. (2020). Structure of the SARS-CoV-2 spike receptor-binding domain bound to ACE2. Nature.

8. Shang, J. et al. (2020). Structural basis of receptor recognition by SARS-CoV-2. Nature.

## Docking / Computational Methods

13. Trott, O. & Olson, A.J. (2010). AutoDock Vina. J Comput Chem.

14. Wallace, A.C. et al. (1995). LigPlot: protein–ligand interactions. Protein Eng.

15. Morris, G.M. et al. (2009). AutoDock4 and AutoDockTools. J Comput Chem.

## Experimental / Assay Support

16. Wrapp, D. et al. (2020). Cryo-EM structure of the SARS-CoV-2 spike. Science.

17. Hoffmann, M. et al. (2020). Camostat inhibits SARS-CoV-2 activation. Cell.

18. Glowacka, I. et al. (2011). TMPRSS2 and viral activation. J Virol.

## Multi-target / Drug discovery context

19. Pushpakom, S. et al. (2019). Drug repurposing progress and challenges. Nat Rev Drug Discov.

20. Lionta, E. et al. (2014). Molecular docking in drug discovery. Curr Top Med Chem.

